# CES1 Deficiency is Associated with Metabolic Reprograming and Endothelial Dysfunction in Pulmonary Arterial Hypertension

**DOI:** 10.1101/2025.04.25.650735

**Authors:** Stuti Agarwal, Lyong Heo, Ananya Chakraborty, Ankita Mitra, Lichao Liu, Natasha Auer, Gowri Swaminathan, Flora Huang, Prakash Chelladurai, Eleana Stephanie Guardado, Juan Matos, Anuradha Bankar, Crystal Le, James West, Karthik Suresh, Ramesh Nair, Marlene Rabinovitch, Christophe Morisseau, Bruce D Hammock, Joseph Wu, Zolt Arany, Mark R Nicolls, Vinicio de Jesus Perez

## Abstract

**Background:** Pulmonary arterial hypertension (PAH) is a progressive disease characterized by pulmonary microvascular loss and obliterative remodeling, driven by metabolic reprogramming, oxidative stress, and endothelial dysfunction. While BMPR2 mutations contribute to metabolic shifts in pulmonary microvascular endothelial cells (PMVECs), their low penetrance suggests additional genetic modifiers play a role. A genetic screen of PAH PMVECs identified carboxylesterase 1 (CES1)**—**an endoplasmic reticulum (ER) enzyme involved in lipid metabolism and detoxification—as a candidate regulator of endothelial metabolism and angiogenesis. We hypothesize that CES1 loss promotes endothelial dysfunction via metabolic reprogramming, lipotoxicity, and oxidative stress.

**Methods:** PAH and healthy PMVECs and lung tissues were obtained from transplant donors and commercial sources. CES1 expression was modulated in PMVECs using siRNA knockdown and plasmid overexpression. Mitochondrial and ER function were assessed via confocal microscopy and proteomics. CES1 heterozygous knockout (HET KO) and endothelial-specific knockout (ECKO) mice were exposed to normoxia or hypoxia, with lung tissues analyzed by single-cell RNA sequencing (scRNA-seq) and histopathology.

**Results:** CES1 expression was significantly reduced in PAH PMVECs and vascular lesions. CES1-deficient PMVECs exhibited increased apoptosis, reactive oxygen species (ROS) production, mitochondrial fragmentation, ER stress, and impaired angiogenesis. Confocal imaging and metabolic studies revealed lipid droplet accumulation, reduced fatty acid oxidation, and a glycolytic shift— phenotypes reversed by CES1 restoration. Mechanistically, CES1 transcription was induced by BMPR2 via NRF2 activation, a key regulator of redox and metabolic homeostasis. In vivo, CES1-deficient mice developed severe pulmonary hypertension (PH) under hypoxia, with extensive vascular remodeling, right ventricular dysfunction, and dysregulated angiogenesis and lipid metabolism pathways, as confirmed by lung scRNA-seq.

**Conclusions:** CES1 is essential for pulmonary endothelial homeostasis and serves as a critical modifier of BMPR2 signaling. Given the limited efficacy of current PAH therapies in reversing endothelial dysfunction and small-vessel loss, restoring CES1 expression represents a promising therapeutic strategy.

## INTRODUCTION

Pulmonary arterial hypertension (PAH) is a rare cardiopulmonary disease characterized by vascular remodeling and loss of pulmonary microvessels, leading to right ventricular (RV) failure and death^1^. Pulmonary endothelial cell dysfunction and microangiopathy are pathologic hallmarks of PAH associated with progressive loss and obstructive remodeling of small pulmonary arterioles^2^, and unfortunately cannot be prevented by current PAH therapies. Given that endothelial dysfunction is a key pathological feature of PAH, understanding the mechanisms that regulate the maintenance and repair of pulmonary vessels could help accelerate the discovery of new therapies for PAH.

Studies by our group and others have shown a high prevalence of insulin resistance, hyperglycemia, and hyperlipidemia in non-obese PAH patients associated with worse functional status and higher mortality^3,4^. PAH lung and heart tissues display evidence of lipotoxicity, including pulmonary artery atherosclerosis, oxidized lipid species and ceramide deposits within lung vascular lesions and RV cardiomyocytes^4–6^. These abnormalities are result of metabolic reprogramming when cells switch from mitochondria-based glucose oxidation and fatty acid oxidation (FAO) to glycolysis (i.e., “Warburg effect”)^7–10^. Metabolic reprogramming promotes endothelial dysfunction through the cumulative effect of oxidative stress, endoplasmic reticulum (ER) stress, and mitochondria damage, leading to apoptosis, inflammation, and vessel loss. Thus, understanding the mechanisms of metabolic reprogramming in PMVECs is a top priority for developing novel therapies for PAH.

BMPR2 insufficiency studies on endothelial-specific BMPR2 knockout mice and patient-derived PMVECs have been linked to metabolic reprogramming^11,12^. BMPR2 is a serine/threonine kinase transmembrane receptor that, upon binding BMP ligands, activates several downstream signaling mediators required for PMVEC homeostasis and vascular repair^13^. BMPR2 mutations are the most common genetic cause of hereditary (∼70%) and sporadic (∼11-40%) PAH^14,15^. However, reduced BMPR2 expression and signaling activity is also common in non-mutant PAH patients. Since only ∼20% of BMPR2 mutation carriers develop PAH, genetic modifiers (i.e., “second hits”) are required for PAH development in BMPR2 insufficiency.

Using a genomic screen of PAH PMVECs, we identified carboxylesterase1 (CES1) as a candidate regulating lipid metabolism and a genetic modifier of BMPR2 signaling in PMVECs^16^. CES1 is an ER enzyme that, through its lipase activity, hydrolyzes cholesteryl esters and triacylglycerols (TG) to release free fatty acids (FFA), which are then shuttled to the mitochondria for FAO. CES1 deficiency is associated with lipotoxicity resulting from intracellular lipid and lipid peroxides buildup^17^. While CES1 deficiency contributes to lipid disorders such as atherosclerosis and non-alcoholic steatohepatitis (fatty liver disease or NASH)^18,19^, the role of CES1 in PAH is unknown. Here, we present evidence that CES1 regulates lipid metabolism and redox balance in PMVECs. We show that loss of CES1 leads to lipotoxicity, ROS production, ER and mitochondrial stress which is associated with metabolic reprogramming and endothelial dysfunction in PAH. in PMVECs. We also show that CES1 expression in PMVECs is co-regulated by BMPR2 signaling and epigenetic modifications at the CES1 gene locus. Most importantly, we demonstrate that restoring CES1 expression to baseline values can restore metabolic homeostasis and improve PAH PMVEC function. Taken together, our findings suggest that targeting CES1 could be a novel treatment approach to restore metabolic homeostasis in PAH.

## MATERIAL AND METHODS

Detailed methods can be found in the supplementary material.

### Animals

All experimental protocols used in this study were approved by the Animal Care Committee of Stanford University and adhered to the published guidelines of the National Institutes of Health on the use of laboratory animals. Mice were housed under a 12-hour light and dark cycle with free access to food and water, under pathogen free conditions. Age and sex-matched 2–3-month-old C57BL/6 male mice (n=6-8 animals per group) were used in the study. To study the role of CES1 gene in PAH pathology, we partnered with the NIH Mutant Mouse Resource and Research Center at UC Davis to generate the <u>first</u> CES1d (i.e. mouse isoform of human CES1) KO mouse using CRISPR to remove exon 4 from the gene. The animals used in the study are heterozygous CES1 KO that is they carried mutation only on one allele. We also used endothelial specific CES1 conditional knockout mice (hereafter referred to as CES1 ECKO mice) obtained by crossing CES1 flox/+ mice and VE-Cadherin-CreERT2 mice for at least 5 generations. Ear clip-based genotyping of the progeny was used to identify littermate controls and CES1 ECKO mice. Mice were treated with 2mg of tamoxifen (20 mg/mL dissolved in corn oil) for 3 consecutive days to induce the CES1 knockout. The wild type (WT) mice with no mutations on any of the alleles were taken as controls.

### Cell Culture

Healthy control and PAH PMVECs were obtained from the Pulmonary Hypertension Breakthrough Initiative (PHBI), with additional healthy donor cells obtained from PromoCell (Sigma-Millipore, USA). Cells were grown in endothelial cell medium (ECM) supplemented with 5% fetal bovine serum (FBS) and growth factors (VascuLife, USA) on fibronectin-coated cell culture dishes. For experiments, PMVECs were plated and grown until 70-80% confluency and then starved using ECM containing 0.5% FBS. Donor clinical information can be found in **Supplementary Table 1**.

### Statistical Analysis

All statistical analyses were performed using Prism 10 software (GraphPad). The minimum level of p< 0.05 was considered significant and reported to the graphs. All experiments were repeated at least three times (independent replicates); n represents a pool across these experimental replicates. Data are presented as means ± SEM. Statistical comparison were performed using the unpaired Student t-test or Mann-Whitney for non-parametric data. Comparison among ≥3 groups was performed using ordinary 1-way ANOVA or 2-way ANOVA with Dunnet’s/Tukey’s post hoc test if the data followed a normal distribution, otherwise non-parametric Kruskal-Wallis test with Dunn post hoc analysis was used. In case of multiple group analyses, all pairwise comparisons were tested; however, only those which appeared to be relevant for the interpretation of the data are presented. Animal experiments were performed with littermates as controls. The choice of animal number was based on power calculations resulting in a power of >0.9 for a SD of 0.2 to find a difference of 0.4.

## RESULTS

### CES1 expression is reduced in PAH PMVECs and lung vascular lesions

PMVEC metabolic reprogramming and oxidative stress is a well-described phenomenon in PAH^10,20^, but the mechanisms responsible are incompletely understood. Previous analysis of RNA sequencing^21^ data by our group for candidate genes linked to metabolism and redox biology led to the identification of CES1 as a significantly <u>downregulated</u> gene in PAH PMVECs. We observed that CES1 expression was significantly reduced in PAH PMVECs at both mRNA (**Fig. 1A**) and protein (**Fig. 1B**) levels. We also demonstrated that CES1 was expressed predominantly in endothelial layer in healthy lung micro vessels, while CES1 expression was significantly reduced within PAH lesions (**Fig. 1C**).

**Figure 1.**
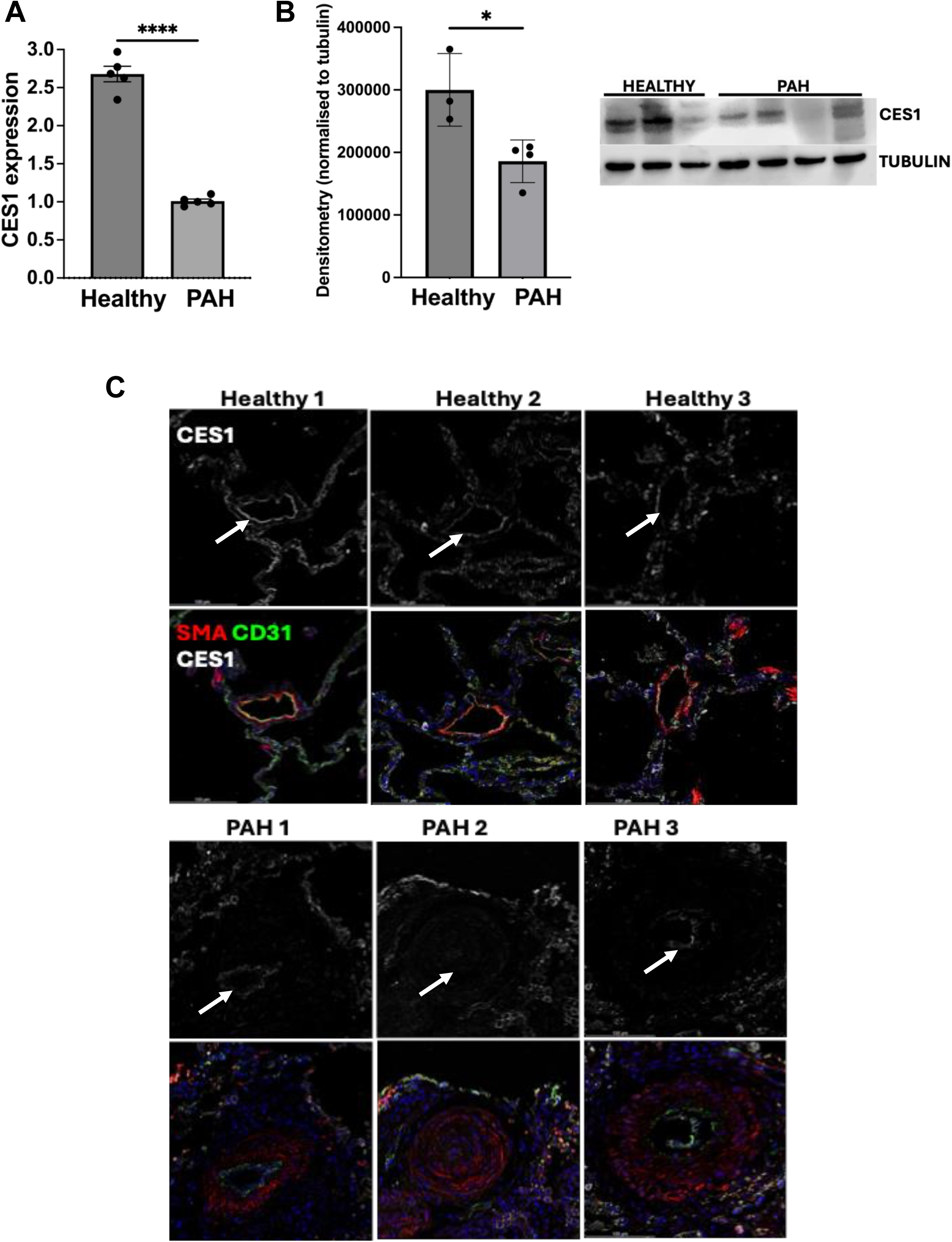
CES1 expression is reduced in PAH PMVECs and lung vascular lesions. A) qPCR, and B) WB analysis for CES1 expression in healthy and PAH HPMVECs *p<0.05, **p<0.01 vs Healthy donors (Mann Whitney test), C) Confocal images showing CD31 (green) and CES1 (gray) staining in human lung cryosections from healthy and PAH patients (Scale-100µm).

### CES1 deficiency causes PMVEC dysfunction and oxidative stress

Next, we determined if loss of CES1 in endothelial cells in PAH is correlated with endothelial dysfunction characterized by increased susceptibility to apoptosis, and loss of pro-angiogenic capacity. Healthy PMVECs were transfected with a CES1-specific siRNA (siCES1) for 72 hours. Compared to controls, siCES1 PMVECs exhibited increased caspase 3/7 activation upon starvation (i.e., increased apoptosis) (**Fig. 2A**). In a Matrigel assay, siCES1 PMVECs formed fewer vascular tubes vs. controls (**Fig. 2B**).

**Figure 2.**
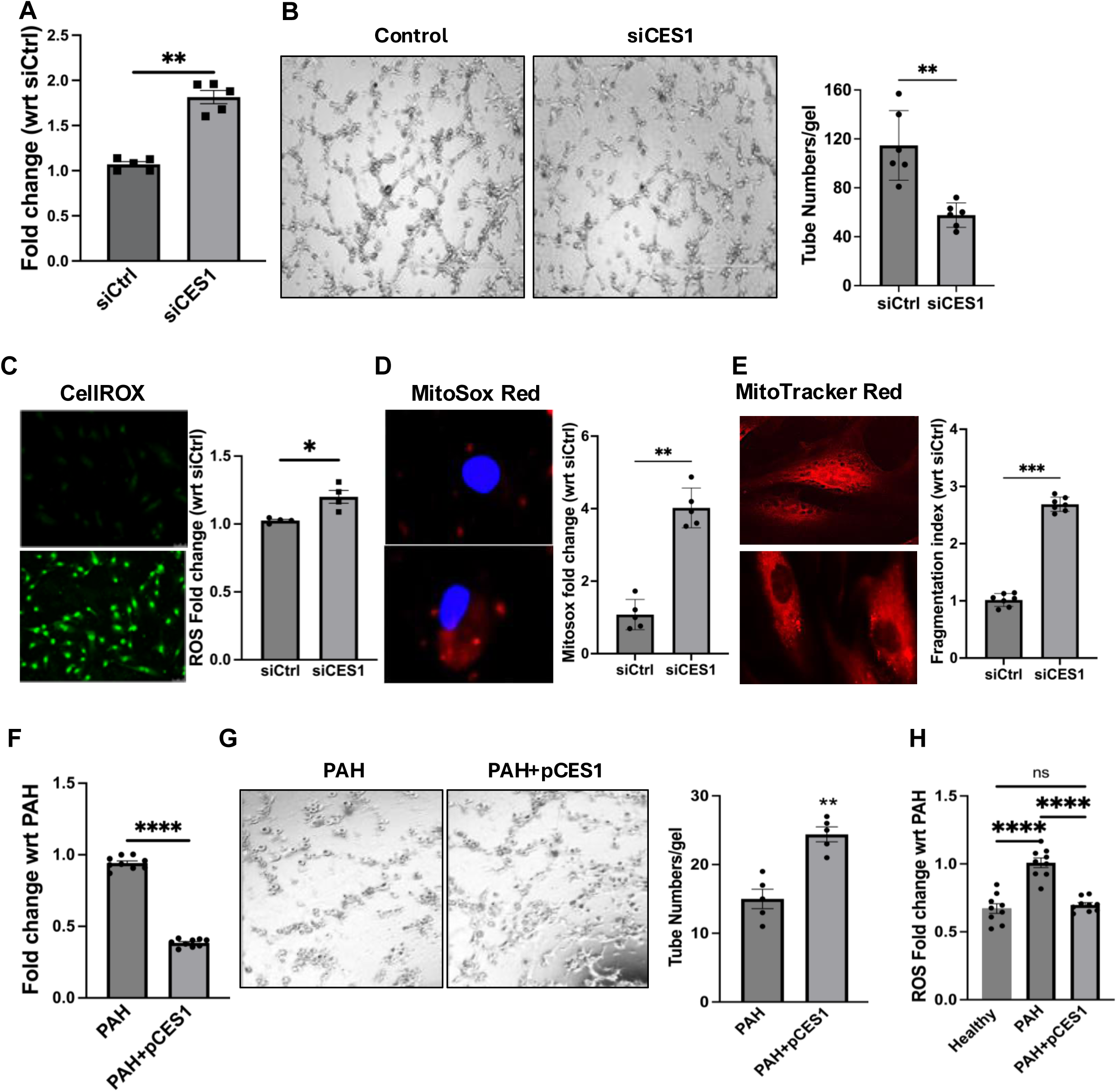
CES1 deficiency leads to endothelial dysfunction and induces oxidative stress. A) Caspase-3 Glo assay showing apoptosis levels, B) Matrigel tube formation assay (Scale-100µm) and C) Total ROS fluorometry in control and siCES1 transfected human PMVECs. D) MitoTracker Red stain, fragmentation index and E) MitoSOX stain, fluorometry measurement of control (top) and siCES1 (bottom) PMVECs. Scale-100µm. F) Caspase-3 Glo assay for apoptosis, G) Matrigel tube formation assay (Scale-100µm) PAH HPMVECs transfected with vector or pCES1. *P<0.05, **P<0.01, ****P<0.0001, Mann Whitney test. H) Total ROS fluorometry of PAH PMVECs transfected with vector or pCES1 and healthy PMVECs. ***P<0.001, ****P<0.0001, one-way ANOVA.

Previous studies have shown that increased oxidative stress is a known feature of endothelial dysfunction and mitochondrial damage in PAH^20,22–26^. Given the involvement of CES1 in redox and metabolic homeostasis^27^, we hypothesized that CES1 knockdown would result in increased ROS production and mitochondrial stress. Compared to controls, siCES1 PMVECs demonstrated significantly higher total cell ROS (**Fig. 2C**)^28^. Mitochondrial superoxide production and increased fragmentation (a marker of fission) are two common markers of mitochondrial dysfunction^20,29^. Compared to controls, mitochondria-specific ROS was greater in siCES1 PMVECs (**Fig. 2D**). While filamentous mitochondria were seen in control PMVECs, siCES1 PMVECs mitochondria were fragmented and distributed haphazardly across the cytoplasm (**Fig. 2E**).

Restoring CES1 in PAH PMVECs transfected with a CES1 expression construct (pCES1) demonstrated significantly lower apoptosis than PAH PMVECs transfected with an empty vector (**Fig. 2F**) as well as increased tube numbers and network formation (**Fig. 2G**). Furthermore, as compared to healthy PMVECs, PAH PMVECs exhibited <u>higher</u> levels of total ROS, while PAH PMVECs transfected with pCES1 demonstrated significantly reduced total ROS under the same conditions (**Fig. 2H**). In conclusion, the results indicate that CES1 is required for PMVECs homeostasis, and its loss can increase susceptibility to apoptosis, oxidative stress and reduced angiogenic capacity.

### CES1 deficiency is associated with ER stress and lipotoxicity

Oxidative stress can result from increased ROS production by mitochondria and ER, two organelles that interact through lipid metabolism. Given the location of CES1 in the ER, we sought to determine whether siCES1 exhibit ER stress in addition to mitochondria dysfunction. Compared to controls, siCES1 PMVECs showed significant increase in ER stress markers (PERK, IRE-1a, CHOP, BiP)^30^ (**Fig. 3Ai and ii**). Given the evidence of mitochondrial fragmentation, siCES1 PMVECs demonstrated reduced OPA-1/mitofusin 1 expression and increased Drp1, favoring mitochondrial fission dynamics (**Fig. 3Bi and ii**). We also conducted transmission electron microscopy (TEM) in siCES1 PMVECs which also showed evidence of distended mitochondria and ER cisternae (**Fig. 3C**), further emphasizing the presence of mitochondrial and ER stress in siCES1 cells.

**Figure 3.**
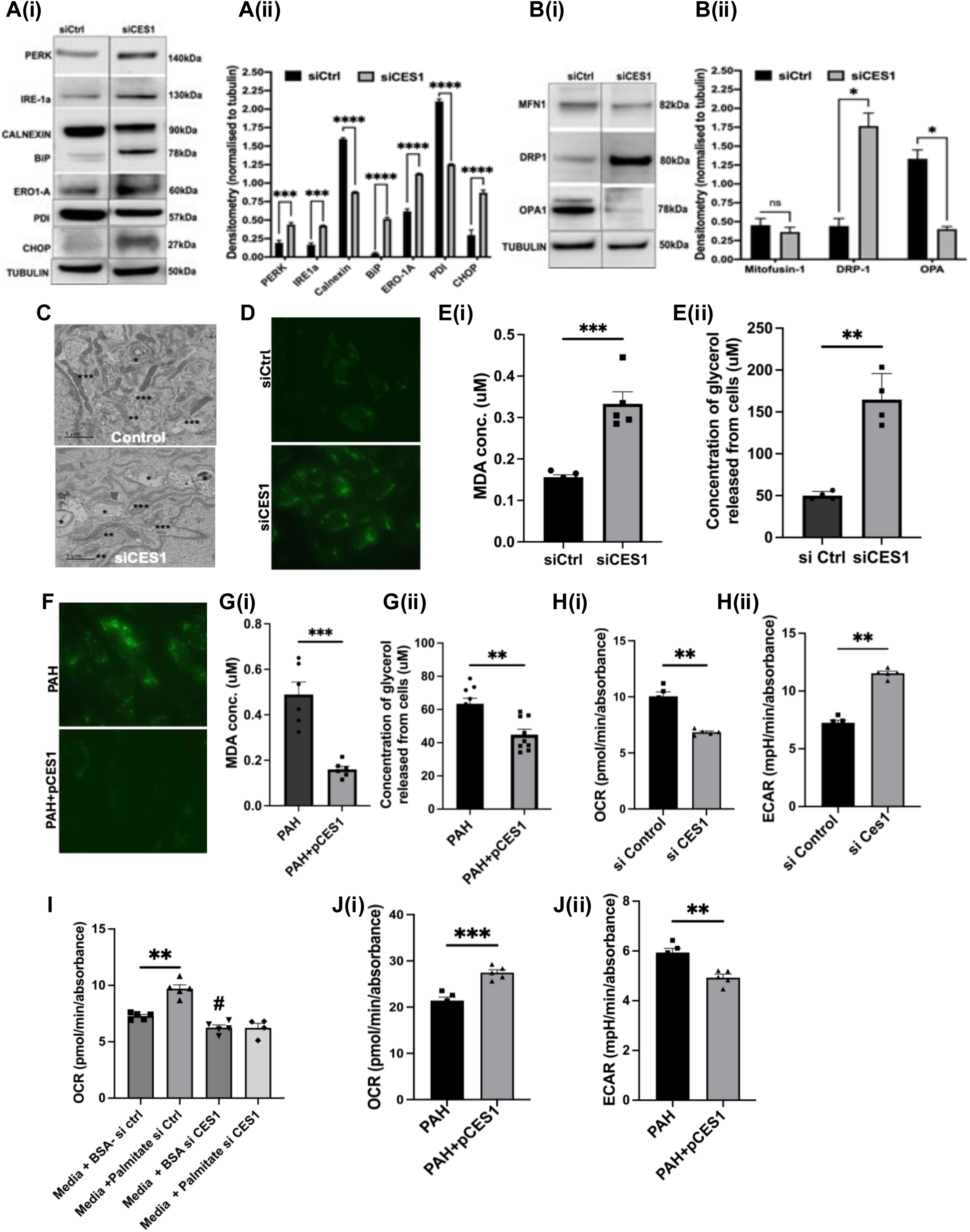
CES1 deficiency causes lipotoxicity, ER/mitochondrial stress and metabolic reprograming. WB and Densitometry of A) ER stress markers and B) mitochondrial fusion and fission markers respectively in whole cell lysates obtained from siCtrl and siCES1 transfected cells. *p<0.05, ***p<0.001, ****p<0.0001, 2-way ANOVA). C) TEM (Transmission Electron Microscopy) demonstrating lipid droplets (*), ER (**), and mitochondria (***) in control and siCES1 PMVECs. D) LipidSpot 488 staining in control and siCES1 PMVECs (green dots represent cytoplasmic lipid droplets, Scale-100µm). E) (i) Lipid peroxidation assay showing Malondialdehyde byproduct formation and (ii) Triglyceride-Glo assay showing quantification of intracellular glycerol released from cells in control and siCES1 PMVECs. *P<0.05, **P<0.01, ***P<0.001, Mann Whitney test. F) LipidSpot 488 staining, G(i) Lipid peroxidation assay and (ii) Triglyceride-Glo assay in PAH PMVECs transfected with vector or pCES1, *P<0.05, **P<0.01, ***P<0.001, Mann Whitney test. H) Graph showing (i) oxygen consumption rate (OCR) and (ii) Extracellular acidification rate (ECAR), and I) OCR by mitochondria upon supplementation of palmitate as substrate for oxidation in control and siCES1 PMVECs. J) Graph showing (i) OCR and (ii) ECAR in PAH PMVECs with and without CES1 overexpression. All experiments were performed in the Seahorse XF96 machine using the manufacture protocol and assays. *p<0.05, **P<0.01, ***P<0.001, Mann Whitney test and one-way ANOVA.

An intriguing finding from the TEM image analysis was the presence of multiple lipid droplets in the cytoplasm of siCES1 PMVECs. LipidSpot and Oil Red O staining confirmed accumulation of cytoplasmic large lipid droplets in both siCES1 (**Fig. 3D**) and PAH PMVECs (**Fig. S1A**). Normally, lipid droplets prevent lipotoxicity from intracellular buildup of triglycerides; however, the accumulation of lipid droplets can also become pathogenic and form lipid peroxides which damage cellular organelles. Lipotoxicity from buildup of lipid peroxides is a common pathogenic feature of PAH associated with vascular remodeling, RV failure and metabolic dysregulation ^6,31–33^. Since CES1 is a known lipase that regulates the release of FFAs from triglycerides, we hypothesized that CES1 deficiency triggers oxidative stress and ER/mitochondria dysfunction through lipid peroxidation. To test for lipid peroxidation, we measured production of malondialdehyde (MDA), a byproduct of lipid peroxidation. Compared to controls, both siCES1 (**Fig. 3Ei**) and PAH (**Fig. S1B)** demonstrated significantly higher cytoplasmic MDA concentration. We also measured triglyceride levels using the Triglyceride-Glo assay and found that triglyceride concentration was also increased in both siCES1 (**Fig. 3Eii**) and PAH PMVECs **(Fig. S1C),** leading us to conclude that triglyceride metabolism was reduced in these cells. Notably, transfection of pCES1 in PAH PMVECs resulted in a significant reduction in cytoplasmic lipid droplet content (**Fig. 3F**), MDA (**Fig. 3Gi),** and triglyceride content (**Fig. 3Gii).**

### CES1 deficiency reduces fatty acid oxidation and promotes glycolysis of PMVECs

Given the lipid droplet accumulation and mitochondrial damage, we hypothesized that this could affect oxidative metabolism in siCES1 PMVECs. Metabolic reprogramming in PAH is associated with an increase in glycolysis, thus, cytoplasmic and mitochondrial glucose oxidation were measured by Glycolytic Rate assay and MitoStress Test assay respectively on Seahorse XF platform. We found significant decrease in Oxygen Consumption Rate (OCR) and concomitant increase in glycolysis were obtained, as evidenced by an increase in Extracellular Acidification Rate (ECAR) in siCES1 (**Fig. 3Hi & ii**) and PAH (**Fig. S1Di & ii**) PMVECs. We measured FAO using the Seahorse Palmitate Oxidation assay and found a significant reduction in OCR for palmitate oxidation (an FFA substrate for FAO) in the siCES1 PMVECs compared to controls (**Fig. 3I**). Finally, we confirmed that CES1 overexpression in PAH PMVECs improved metabolic activity as evidenced by an increase in OCR (**Fig. 3Ji)** and a reduction in ECAR (**Fig. 3Jii**). Taken together, our studies provide evidence that CES1 deficiency can induce metabolic reprogramming in PMVECs.

### RNA sequencing of CES1 deficient PMVECs reveal modulation in lipid metabolism and angiogenic pathways

To identify the mechanisms by which CES1 deficiency results in PMVEC and metabolic dysfunction, we conducted bulk RNA-seq of control and siCES1 PMVECs. Heat map (**Fig. 4A**) and volcano plot (**Fig. S2A**) was generated for overall top 10 upregulated and downregulated genes in siCES1 PMVECs. Some of the most important genes that were modulated in siCES1 PMVECs were PEG10 (paternally expressed gene10) that regulates cell growth and differentiation, PTGS2 (prostaglandin-endoperoxide synthase 2) responsible for fatty acid metabolism and cell proliferation, IL1B (interleukin-1 beta) and IL6 (interleukin 6), pro-inflammatory cytokines that are known to be modulated in PAH. We then performed gene set enrichment analysis (GSEA) analysis for gene ontology (GO): Biological Process (BP) in which top upregulated processes included regulation of cell growth and response to ER stress while top downregulated processes included lipid and fatty acid metabolic pathways (**Fig. 4B**).

**Figure 4.**
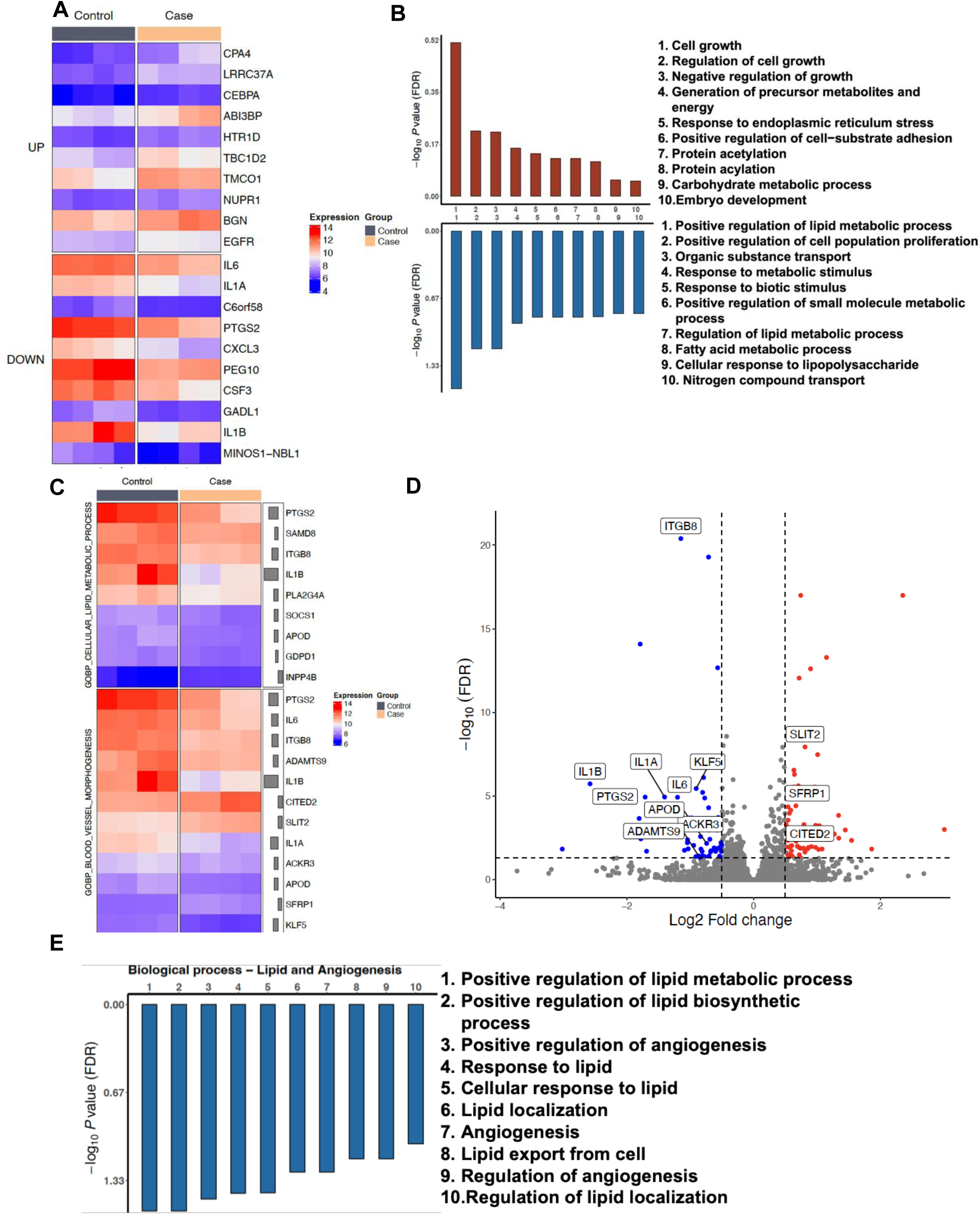
RNAseq of CES1 deficient PMVECs reveals modulation of metabolic and angiogenic pathways. Heatmap analysis of A) top 10 upregulated and down regulated genes (blue to red defines low to high log2fold change) and B) Gene set enrichment analysis (GSEA) for GO: Biological Process (BP) showing top 10 up-regulated (red bars) and down-regulated (blue bars) biological pathways in siCES1 PMVECs as compared to control PMVECs. All terms are significantly enriched using adjusted p value < 0.05 on normalized enrichment scores (NES) that were computed by GSEA on fold change-ranked genes. C) Heatmap analysis and D) Volcano plot showing highly up and down regulated genes in gene ontology (GO) pathways for lipid metabolism and angiogenesis biological processes in siCES1 PMVECs as compared to controls, blue to red defines low to high log2fold change. E) Gene set enrichment analysis (GSEA) for biological processes (BP)-cellular lipid metabolism and blood vessel morphogenesis showing top 10 down-regulated (blue bars) biological pathways in siCES1 PMVECs as compared to control PMVECs. All terms are significantly enriched using adjusted p value < 0.05 on normalized enrichment scores (NES) that were computed by GSEA on fold change-ranked genes.

Depending on the overall DE analysis and experimental evidence of lipid accumulation and reduced angiogenesis in CES1 deficient PMVECs, two GO terms-GO:0044255 (cellular lipid metabolic process) and GO:0001525 (blood vessel morphogenesis/ angiogenesis) were chosen to look at differential gene expression for specific BP of interest. Heatmap **(Fig. 4C)** and volcano plot **(Fig. 4D)** were generated for differential gene expression. These included ITGB8 (integrin beta 8) involved in cell-cell adhesion and APOD (Apolipoprotein D) which plays role in lipid metabolism. Selective GSEA for two GO:BP terms-lipid metabolism and angiogenesis interestingly only revealed downregulated processes like cell response to lipid, lipid transport and regulation of angiogenesis as shown in **Fig. 4E**.

### CES1 is a downstream target of BMPR2-NRF2 signaling

It is known that BMPR2 insufficiency in PMVECs is associated with mitochondrial dysfunction, lipid peroxidation, metabolic reprogramming and oxidative stress^20,26,31,32^. Given the similarity in phenotypes observed with CES1 deficiency, we explored if BMPR2 and CES1 are part of a common signaling axis. CES1 gene and protein expression were measured in PMVECs transfected with control or BMPR2-specific siRNA (siBMPR2). Interestingly, compared to control, siBMPR2 exhibited significantly reduced CES1 expression at both mRNA **(Fig. 5A)** and protein levels **(Fig. 5B)** at 48 hours after transfection. Next, apoptosis rates were compared in siCES1 vs. siBMPR2 PMVECs in starvation media for 24hrs. in the presence or absence of BMP-9 (10ng/ml)^13,34^. As expected, siBMPR2 and siCES1 PMVECs exhibited increased apoptosis in media (**Fig. 5C**). With BMP-9, there was a significant reduction in apoptosis in controls but not much in siBMPR2 PMVECs. Intriguingly, apoptosis levels did not change even in siCES1 PMVECs, suggesting that CES1 is required for the pro-survival activity of BMPR2 in PMVECs (**Fig. 5C**).

**Figure 5.**
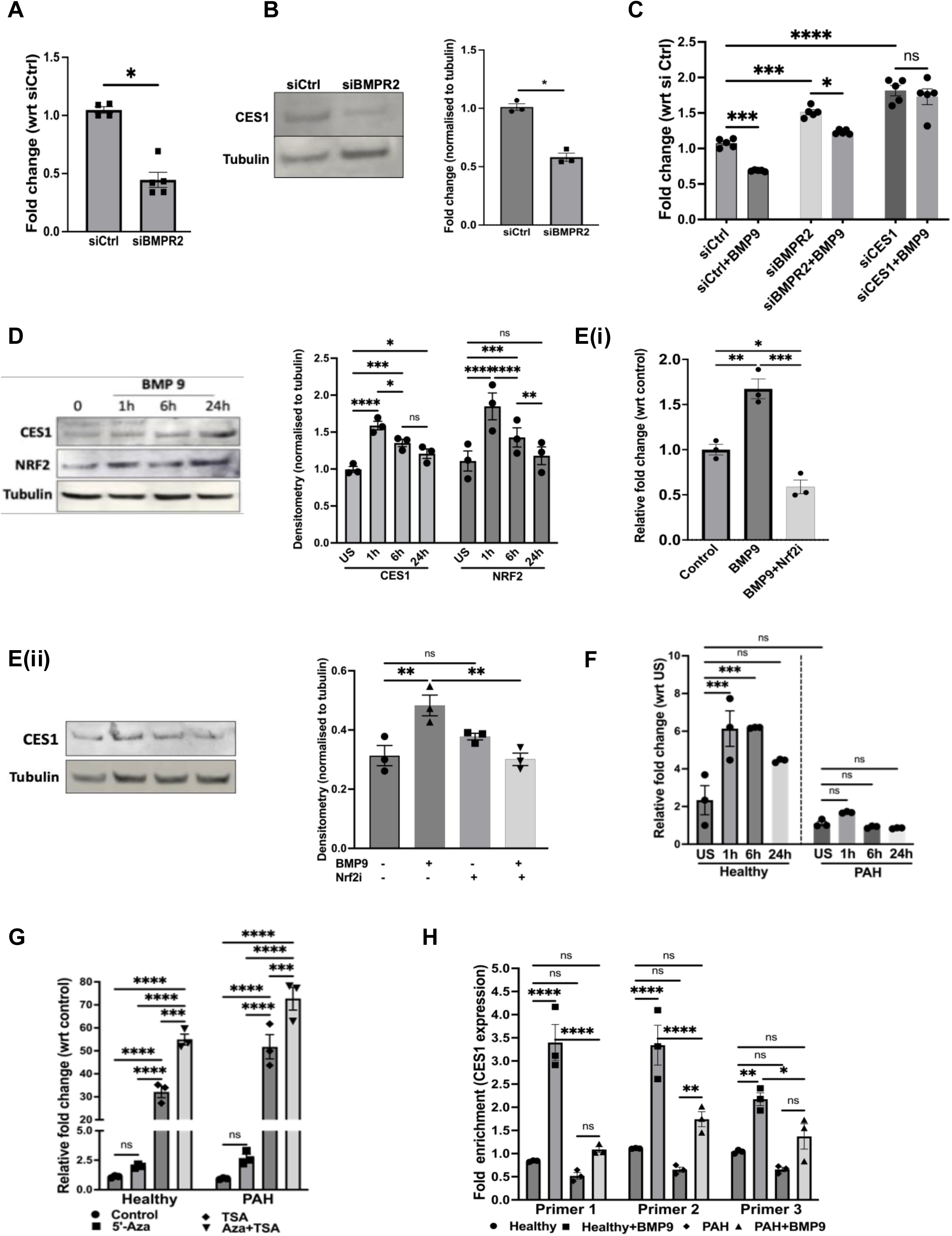
CES1 expression in PMVECs is linked to BMPR2-NRF2 signaling and controlled by epigenetic mechanisms. A) Quantitative PCR and B) WB of CES1 in control and siBMPR2 PMVECs. *P<0.5, **P<001, Mann Whitney test, C) Caspase 3/7 assay of siBMPR2 and siCES1 PMVECs in serum free media +/− BMP-9 for 24 hours. D) WB of NRF2 and CES1 in HPMVECs stimulated with BMP9 for the timepoints 0 h (unstimulated control), 1 h, 6 h, and 24 h; E(i) mRNA expression and (ii) protein expression of CES1 in cell lysates of HPMVECs stimulated with BMP9 in siNRF2 PMVECs. F) qPCR for *CES1* in Healthy donor and PAH PMVECs stimulated with BMP-9 for 24 hours, G) qPCR for *CES1* in healthy and PAH PMVECs stimulated with BMP-9 for 6 hours and exposed to 5-AZA and TSA, H) qPCR to show *CES1* chromatin bound to NRF2 binding motif on *CES1* promotor sites. *p<0.05, **p<0.01, ***p<0.001, ****p<0.0001, two-way ANOVA.

BMPR2 activation can trigger gene expression changes through canonical and non-canonical signaling pathways^35,36^. In the canonical pathway, BMPR2 acts by triggering the phosphorylation of SMADs transcription factor. Non-canonically, BMPR2 activates MAP kinases (Akt, JNK, p38)^36^, and can directly activate transcription factors such as PPARγ and NRF2 involved in cell metabolism and PAH^37–39^.

Contrary to our expectations, screening the CES1 gene locus revealed no predicted canonical Smad-binding motifs. However, there is evidence of multiple NRF2 binding sites within the CES1 promotor and enhancer sequences (**Fig. S3A**). NRF2 is a highly conserved transcription factor that binds to <u>A</u>ntioxidant <u>R</u>esponse <u>E</u>lements (ARE) consensus sequences in the promoters and enhancers of antioxidant genes involved in the oxidative stress compensatory response^39,40^, mitochondrial biogenesis (via TFAM and PGC-1α) and respiration^41,42^. To test whether BMPR2 can activate NRF2, healthy PMVECs were stimulated with BMP-9 for 24hrs. followed by NRF2 estimation in lysates. BMP-9 stimulation increased NRF2 starting at 1 hour, and levels remained elevated at 24 hours; interestingly, this dynamic increase preceded a *rise* in CES1 protein, reaching its peak at 24 hours (**Fig. 5D**). To test whether NRF2 activation is required for BMP-9 induced CES1 upregulation, cells were transfected with siNRF2 followed by BMP-9 stimulation for 24hrs. Compared to the control, both CES1 mRNA and protein expression remained neutral in siNRF2 PMVECs stimulated with BMP-9 (**Fig. 5Ei and ii**). BMP-9 restores BMP signaling in PAH by increasing BMPR2 expression^38^, hence, we hypothesized that BMP-9 stimulation would increase CES1 expression in PAH PMVECs. Compared to the donor control, BMP-9 induced only a mild increase in CES1 expression in PAH PMVECs suggesting an alternative mechanism for CES1 regulation (**Fig. 5F**).

### CES1 expression in PAH PMVECs is repressed by epigenetic mechanisms

Epigenetic repression plays a key role in PAH pathobiology and that pharmacotherapies that reverse CpG island methylation or inhibit HDAC activity can improve vascular remodeling and RV function in several animal models of PAH^43–46^. *In silico* analysis of the *CES1* gene locus demonstrates the presence of DNA methylation sites and multiple histone marks responsible for gene silencing (**Fig. S3B**). To determine whether patterns of DNA methylation at the NRF2 binding sites differ between PAH vs. donor PMVECs, we used a published DNA Methyl-Seq database to compare global patterns of *CES1* gene methylation in healthy vs. PAH PMVECs derived from patients with idiopathic PAH^46^. Compared to healthy donors (N=7), idiopathic (N=4) PAH PMVECs demonstrated a mild but significant increase in the methylation at CpG island adjacent to the NRF2 binding sites (**Fig. S3C**). To determine acetylation status, we used a ChIP-Seq dataset from healthy vs. PAH PMVECs^47^ to compare the distribution of H3K27Ac (i.e., histone acetylation) marks across the NRF2 binding sites of the CES1 gene. Compared to healthy donors (N=3), H3K27Ac covering the promoter/enhancer was reduced in PAH PMVECs (N=3), suggesting lower chromatin accessibility at CES1 promoter in PAH (**Fig. S3D**).

Based on our data, we hypothesized that <u>reversal</u> of DNA methylation and/or inhibition of histone deacetylation would increase *CES1* expression in PAH PMVECs at baseline and in response to BMP-9. We treated PMVECs with 5-azacytidine (5-AZA, a DNMT inhibitor, 10nM) and trichostatin-A (TSA, a class I/II HDAC inhibitor, 100ng/ml) either alone or together for 24hrs, followed by qPCR for *CES1*. Compared to 5-AZA alone, TSA treatment of PAH PMVECs resulted in greater recovery of *CES1* expression in PAH PMVECs. Despite activation of BMP9/BMPR2 signaling, combined inhibition of DNA methylation and HDAC inhibition was required to significantly enhance *CES1* expression in both healthy and PAH PMVECs demonstrating the importance of chromatin accessibility in determining CES1 transcription (**Fig. 5G**). To assess if BMP-9 modulates NRF2 occupancy at CES1 promoter, we then performed chromatin immunoprecipitation (ChIP) to validate and quantify NRF2 occupancy at the NRF2 binding motifs within the CES1 promotor region. PMVECs from PAH patients showed lower NRF2 occupancy at the CES1 promoter when compared to healthy PMVECs. BMP-9 stimulation significantly enhanced NRF2 occupancy at the *CES1* promoter in both healthy and PAH PMVECs **(Fig. 5H)**, which correlated with the BMP9-induced increase in *CES1* transcription observed upon inhibition of epigenetic repression as shown in **Fig. 5G**.

Thus, we conclude that **(1)** CES1 silencing in PAH PMVECs is caused predominantly by epigenetic repression, and that (**2)** the combination of BMPR2 insufficiency <u>and</u> epigenetic repression can act as a “double hit,” leading to severe CES1 deficiency and endothelial dysfunction (**Fig. S3E**).

### Global heterozygous and endothelial-specific CES1 deficient mice develop severe pulmonary hypertension and vascular remodeling in hypoxia

CES1 is a critical regulator of lung endothelial homeostasis, but whether CES1 deficiency can produce and/or exacerbate pulmonary hypertension *in vivo* remains unknown. To determine the impact of CES1 deficiency, a CES1 KO was generated via CRISPR-based deletion of exon 4 of the *Ces1d* gene, the murine isoform of CES1. CES1 heterozygous mice (CES1 KO) was chosen since homozygous mice die in-utero. Heterozygous CES1 KO mice develop normally and exhibit no discernible phenotype. Endothelial specific CES1 conditional KO mice (CES1 ECKO) was also generated to test the development of PH phenotype upon hypoxia exposure. While there were mildly significant differences between wild type (WT) and CES1 KO/ CES1 ECKO in normoxia, both CES1 KO and CES1 ECKO mice exposed to hypoxia (10% O_2_ for 3 weeks) developed significantly higher right ventricular systolic pressures (RVSP) (**Fig. 6A and S4A**) as well as RV hypertrophy determined by increased RV/LV+Septum weight ratio compared to WT mice (**Fig. 6B and S4B**). This correlated with increased RV area under short axis view on echocardiograph (**Fig. 6C and S4C**). Other echocardiogram parameters suggestive of right ventricle and pulmonary resistance such as RV fractional shortening (RVFS) (**Fig. 6Di and S4Di**) and PAT/PET ratio (**Fig. 6Dii and S4Dii**) were also decreased under hypoxia in both mouse models. However, not much change was seen in left ventricle ejection fraction (LVEF) (**Fig. 6Diii and S4Diii)** and cardiac output (CO) (**Fig. 6Div and S4Div)** suggesting normal left ventricle (LV) function. The CES1 KO and CES1 ECKO hypoxic mice also demonstrated increased vascular remodeling particularly in small arterioles of size less than 50µm (**Fig. 6Ei and ii, S4Ei and ii**). These studies indicate that CES1 deficiency increases the severity of PH associated with hypoxia and acts as a “second hit” in PAH development.

**Figure 6.**
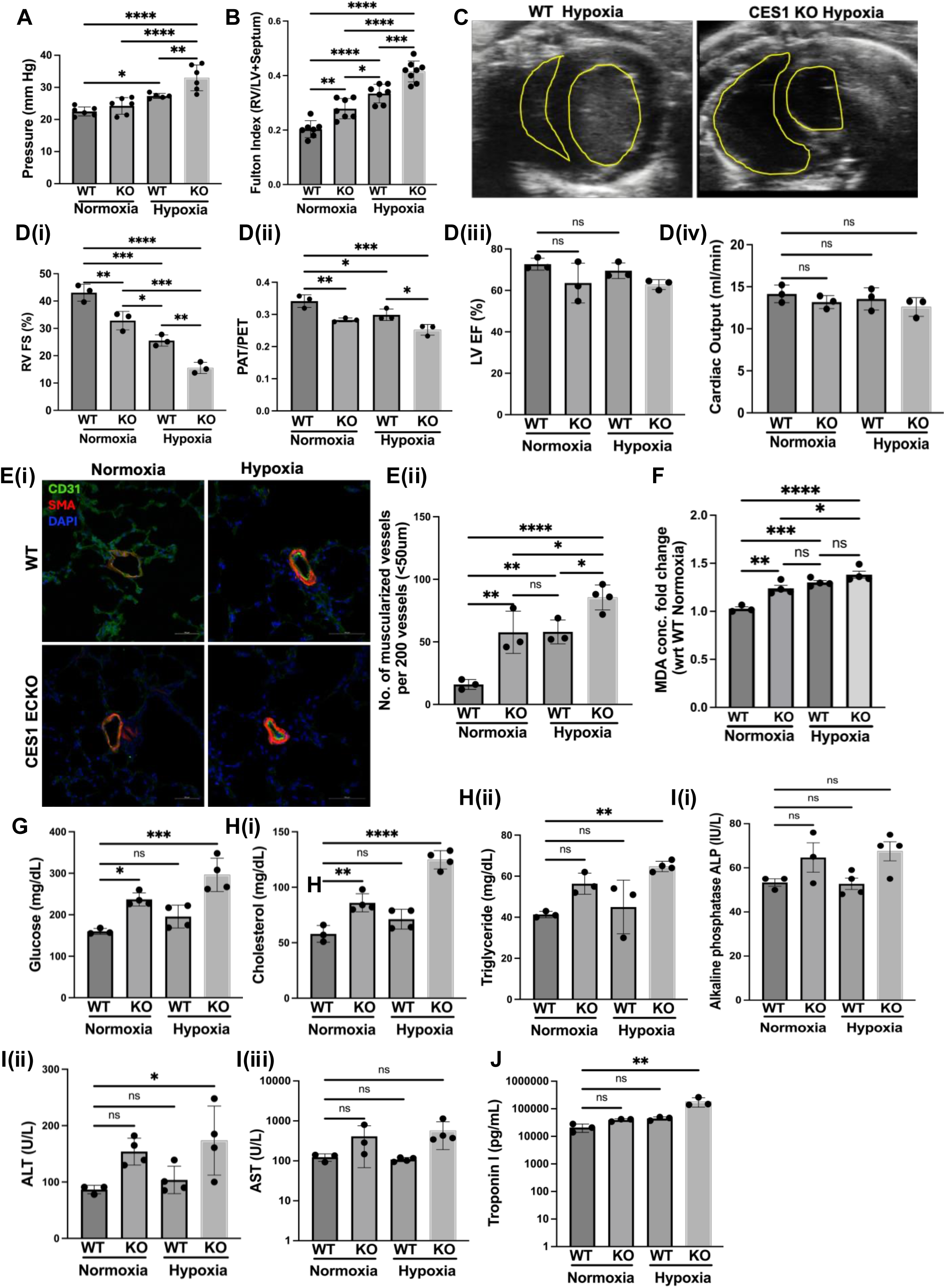
PH phenotype in CES1 HET-KO mice: Hemodynamics and echocardiogram data on age and sex matched WT and CES1-KO mice exposed to normoxia (21% O2) and hypoxia (10% O2). A) Right ventricle systolic pressure (RVSP), B) RV/LV+S (i.e., Fulton) index measurement, C) Comparison of right ventricle area, D(i) Right ventricle Fractional Shortening (RVFS) percentage, (ii) Ratio of Pulmonary Artery Acceleration Time (PAT) and Pulmonary Artery Ejection Time (PET), (iii) Left Ventricle Ejection Fraction (LVEF) percentage and (iv) Cardiac Output in all four mice groups. E) (i) Confocal micrographs showing immunofluorescence staining of lung paraffin-embedded sections with CD31 (green) for endothelial cells and SMA (red) for smooth muscle cells in the pulmonary vessels, DAPI for nuclei, Scale-100µm and (ii) Graphs showing no. of muscularized vessels per 200 vessels of size less than 50µm in WT and CES1 HET KO mice in normoxia and hypoxia. WT and CES1 HET KO mice whole lung tissue lysates were analyzed for formation of lipid peroxides and serum was tested for metabolic panel. Graphs showing F) Lipid peroxidation levels, G) Glucose, H) Lipid panel (i) Cholesterol, (ii) Triglyceride, I) Liver enzymes (i) AST (ii) ALT (iii) ALP and J) Troponin I for cardiac damage in all four mice groups. *p<0.05, **p<0.01, ***p<0.001, ****p<0.0001, one-way ANOVA.

### Heterozygous CES1 knockout mice (CES1 KO) exhibit metabolic abnormalities

Our *in vitro* studies showed that CES1 deficient PMVECs exhibited increased formation of lipid peroxidation products. Compared to WT, CES1 KO mice demonstrated increased lung peroxidation (**Fig. 6F**). Metabolic profiling was conducted in serum from CES1 KO and WT mice. Interestingly, we observed significantly increased glucose (**Fig. 6G)** and lipid (cholesterol and triglyceride) levels (**Fig. 6Hi and ii**) in CES1 KO mice in both normoxic and hypoxic conditions as compared to WT mice. Since CES1 is an important detoxifying enzyme in liver, liver function was checked in all mice groups. Compared to WT mice, all liver enzyme markers-aspartate aminotransferase (AST), alanine aminotransferase (ALT) and alkaline phosphatase (ALP) were in their normal range albeit close to higher range (**Fig. 6I i-iii**). Serum Troponin I were also significantly high in CES1 KO hypoxic mice suggestive of cardiac stress (**Fig. 6J**). Taken together, these findings emphasize that CES1 play a crucial role in maintaining metabolic homeostasis in cells as well as *in vivo*.

### Single-cell RNA-sequencing of CES1 KO lungs reveal alterations in lipid metabolism and lipid transport pathways

The cells from lungs of all 4 mice groups were clustered based on their expression profile, and cell types were annotated based on established cell markers available on LungMap (https://lungmap.net/), CellMarker, The Human Protein Atlas (http://www.proteinatlas.org). A total of 34 clusters were identified, corresponding to six major cell groups: epithelial, stromal, endothelial, myeloid, lymphoid, and mesothelial cells (**Fig. S5A and B**). The overall differentially expressed genes in 4 different groups of mice were identified based on log_2_ fold change and p-value as shown in heat map and volcano plot (**Fig. S5Ci and ii**).

Out of all cell types, we identified five distinct endothelial cell (EC) subset clusters based on their expression profiles of cellular markers corresponding to each EC type as shown in **Fig. S5D**. These included arterial, venous, capillary and lymphatic endothelial cells. Capillary cells formed two distinct clusters gCap and aCap corresponding to capillary cell phenotypes recently characterized by Gillich et al^48^ and their distribution was found to be modulated and present in highest percentage (**Fig. 7A and B**). Interestingly, single-cell RNA-seq analysis revealed a prominent decrease in the proportion of aCap cells followed by arterial endothelial cells in hypoxic CES1 HET KO mice, while other endothelial subsets remained relatively unchanged. The EC specific differentially expressed genes were identified based on log_2_ fold change and p-value in aCap, gCap and all endothelial cells combined as shown in heat map (**Fig. 7C**) and volcano plot (**Fig. S5E**).

**Figure 7:**
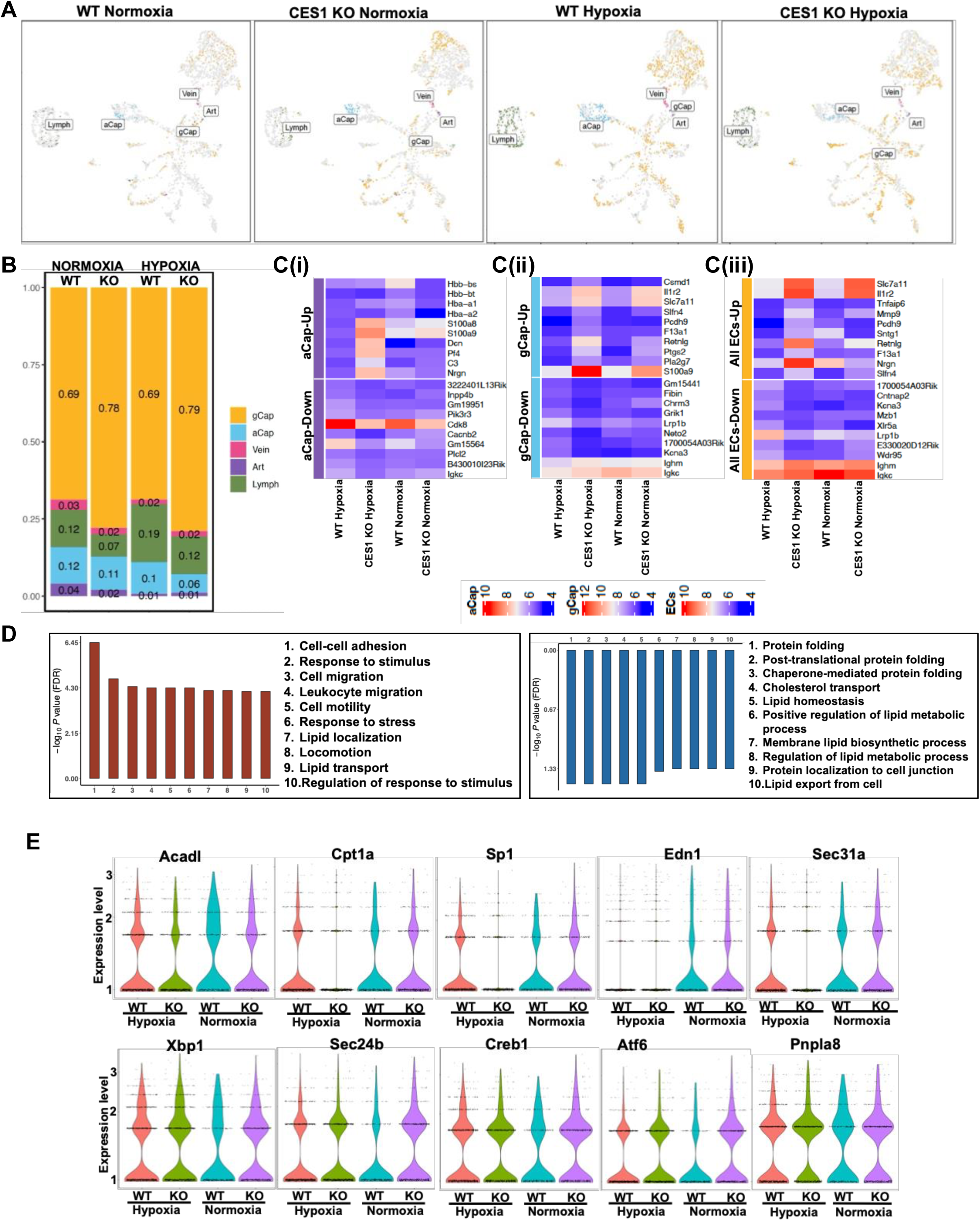
Cellular composition of lung endothelium and gene/pathway enrichment analysis in WT and CES1 HET-KO mice. All data is representative of all endothelial subsets/ clusters in all 4 animal groups-WT and CES1 HET-KO in normoxia and hypoxia. A) UMAP plot of endothelial cell (EC) clusters that were identified, B) graph showing proportion distribution of five clusters of endothelial cells, C) Heatmap of differential expressed genes (DEGs) in (i) aCap cells, (ii) gCap cells and (iii) all ECs combined using pseudo bulk DE analysis (blue to red-low to high log2fold change). D) Top 10 up-regulated (red bars) and down-regulated (blue bars) biological pathways enriched in ECs after gene set enrichment analysis (GSEA) using gene ontology (GO) analysis for biological process (BP) in CES1 HET KO hypoxic mice. All terms are significantly enriched using adjusted p value < 0.05 on normalized enrichment scores (NES) that were computed by GSEA on fold change-ranked genes. E) Expression of genes in all ECs (combined) associated with various lipid biological processes-Acadl and Cpt1a (carnitine metabolic process), Sp1, Sec31a, Sec24b, Atf6 and Creb1(SREBP signaling process) and Edn1, Xbp1, Pnpla8 (fatty acid biosynthetic process) in all 4 mice groups.

Based on the gene sets obtained from pseudobulk differential expression (DE) analysis, GSEA for GO:BP was performed and biological processes were enriched based on the significantly differentially expressed genes in our data set. It was observed that biological processes related to cell death, adhesion and migration were upregulated while pathways associated with lipid localization and metabolism were downregulated in CES1 KO hypoxia group (**Fig. S5F**). Like overall cell cluster analysis, GSEA was performed on endothelial cell subsets using GO:BP and biological processes related to cell adhesion, migration and lipid transport were upregulated while pathways associated with lipid export and lipid homeostasis were downregulated in CES1 KO hypoxia mice (**Fig. 7D**).

As lipid associated biological process was the most predominantly modulated, we checked the expression of lipid metabolism genes in all ECs (combined) in all 4 different mice groups. In the CES1 KO mice exposed to hypoxia, we found that genes Acadl (acyl-CoA dehydrogenase, long chain) and Cpt1a (carnitine palmitoyl transferase 1A) belonging to carnitine metabolic process essential for breaking down fats in mitochondria for energy production, Sp1 (transcription factor for regulating cell cycle and proliferation) and Sec31a (helps in vesicle release from ER) belonging to sterol response element binding proteins (SREBP) signaling process and Edn1 (endothelin-1) belonging to fatty acid biosynthetic process and responsible for cell proliferation were downregulated in ECs (**Fig. 7E**). Other genes such as Sec24b, cAMP response element binding protein (Creb1) and activating transcription factor (Atf6) responsible for ER-Golgi function and ER stress response and X-box binding protein (Xbp1) and patatin-like phospholipase domain-containing (Pnpla8) responsible for cellular stress response, cell migration/ metastasis and lipid metabolism were elevated in CES1 KO hypoxia group (**Fig. 7E**). Of note, Glutathione peroxidase 4 (Gpx4), elongation of very long chain fatty acid (Elovl5), Lipoprotein lipase (Lpl), Interleukin 1 beta (Il1b) and secretion associated RAS related GTPase (Sar1b) remain unchanged within different animal groups (**Fig. S5G**). Overall, scRNAseq data also hints at the role of CES1 in modulating endothelial cell function and metabolic reprogramming.

## DISCUSSION

Metabolic reprogramming is a key pathogenic mechanism in the development and progression of PAH. Available clinical evidence has identified insulin resistance and lipid dysregulation as biomarkers associated with PAH prognosis and a small clinical trial of dichloroacetate (an activator of mitochondrial metabolism) showed promising albeit inconclusive results^49^. Based on the available data, there is an ongoing interest in identifying molecular mechanisms behind the metabolic reprogramming that could serve as therapeutic targets. In this work, a novel paradigm is provided that positions CES1 as a crucial regulator of PMVEC metabolism that explains the association between BMPR2 insufficiency, lipotoxicity, metabolic reprogramming, and oxidative stress in PAH PMVECs (see model, **Fig. 8**). Our previous studies identified CES1 as a susceptibility gene in PAH associated with methamphetamine abuse (METH-PAH) by facilitating detoxification of METH-induced metabolites. We identified a CES1 variant via whole exosome sequencing associated with reduced enzymatic activity in METH-PAH patients, which was not present in IPAH patients^28^. The current data points at a significant downregulation of CES1 that appears to be secondary to a combination of epigenetic repression and insufficient BMPR2 activity.

**Figure 8:**
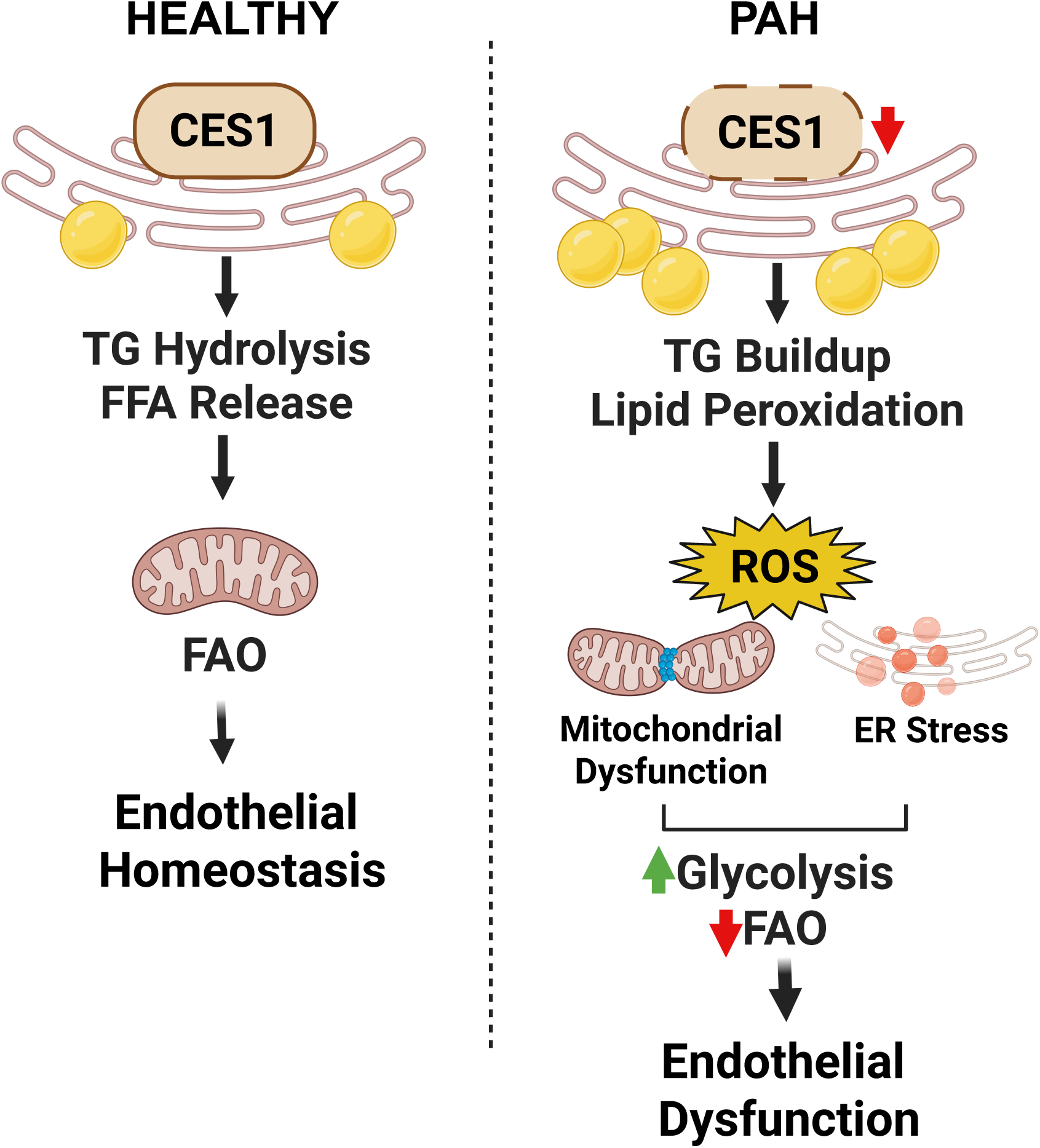
Proposed model describing that CES1 deficiency leads to oxidative stress mediated metabolic reprogramming and endothelial cell dysfunction in PAH.

Carboxylesterases are expressed in many tissues, including the liver, adipose tissue, and endothelial cells. These enzymes play a crucial role in lipid metabolism by catalyzing the hydrolysis of triglycerides, phospholipids, and cholesterol esters. Additionally, CES1 has been shown to play a role in the metabolism of xenobiotics and drugs, and its deficiency may affect the expression of genes involved in drug metabolism and transport. Recent studies have suggested that reduced levels of CES1 activity are associated with endothelial dysfunction, a key event in the development of atherosclerosis and cardiovascular disease^19,27^. Endothelial dysfunction is characterized by impaired endothelium-dependent vasodilation, increased oxidative stress, and inflammation in the endothelial cells (ECs) that line the blood vessels. The exact mechanisms underlying the association between reduced CES1 and endothelial dysfunction are not fully understood. Our study shows for the first time that CES1 is required to maintain metabolic homeostasis through its lipolytic activities which are necessary to maintain a steady supply of FFA for ß-oxidation in the mitochondria. In absence of lipid oxidation to support cell bioenergetics, the cell switches to glycolysis compensate ATP generation and, in doing so, creates a pathological state associated with increased oxidative stress and endothelial dysfunction. One of the major questions raised by our study is how CES1 deficiency generates a state of lipotoxicity through the accumulation of toxic lipid species and causes cell death. This is possible as CES1 is an esterase that breaks down fatty acid esters including cholesterol and triglycerides. Another possibility is that CES1 deficiency may upset the expression and activity of other proteins involved in lipid transport and metabolism which may also contribute to the lipid dysregulation seen in PAH. Evidence of toxic lipid accumulation such as ceramides has been shown in the right ventricle of BMPR2 mutant mice as well as PAH patients^31,32^. Formation of lipid peroxides have been associated with ferroptosis, an-iron dependent apoptosis that leads to metabolic dysfunction in PAH^50^. In future studies, we will explore if lipotoxicity might activate ferroptosis mediated endothelial cell death in CES1 deficiency.

Here, we have focused on BMPR2-NRF2 signaling axis and epigenetics as regulatory mechanisms involved in CES1 expression in PMVECs. However, the regulation of CES1 expression is complex and likely involves multiple transcriptional and post-transcriptional mechanisms. We recognize that BMPR2 can activate other transcription factors such as PPAR-γ and β-catenin that our group and others have shown to be involved in metabolic regulation, CES1 regulation, and lung endothelial homeostasis^20,37,38^. Furthermore, other pulmonary vascular cells affected by CES1 insufficiency might demonstrate cell-specific consequences on their functional status. We plan to expand our research to other cell types and explain how CES1 contributes to vascular remodeling in PAH.

Single cell RNA-seq allows for the identification of different cell types within a tissue and the analysis of their gene expression profiles at a single-cell resolution. In this context, scRNAseq can provide insights into the transcriptional changes that occur in individual cells because of CES1 deficiency. Our RNA-seq data of CES1 deficient PMVECs also revealed modulation of pathways involved in lipid metabolism and transport. Our scRNA-seq revealed that hypoxia exposure in global CES1 KO altered metabolic pathways, particularly those related to lipid metabolism and transport mainly in endothelial cell subtypes. The molecular signature in CES1 KO mice corroborated with increased glucose and lipid levels in the serum from these mice. Our findings suggest that CES1 deficiency can lead to dysregulated lipid metabolism, which may contribute to the development of pulmonary hypertension and other metabolic disorders.

In conclusion, we show a novel mechanism for metabolic reprogramming in PAH based on CES1 subject to regulation by BMPR2 and epigenetic mechanisms. These have implications beyond PAH as it could be applied to other metabolic disorders characterized by lipid dysregulation (e.g., CAD, NASH)^17,19,27^ associated with CES1 deficiency. The discovery of CES1 activators through high throughput screening is a possibility that will be pursued based on our outcomes and should provide exciting opportunities for therapeutic development. With the availability of PAH GWAS and population genetic studies, we will seek to identify and validate potential pathogenic variants of CES1 with direct relevance to clinical PAH which could serve as potential biomarkers and targets for pharmacological and gene editing approaches.

## AUTHOR CONTRIBUTION

S. Agarwal and V.A. de Jesus Perez was responsible for design and performance of experiments, data analysis, interpretation and drafting of the manuscript. All authors contributed to the design, performance and analysis of the studies included in the manuscript. All authors were involved in reviewing and approving the final manuscript.

## Acknowledgements

We acknowledge Cell Sciences Imaging Facility, Stanford University School of Medicine for performing TEM on our samples and Genome Sequencing Service Center, Stanford University for performing mRNA sequencing. We also acknowledge bioinformatics services and computing resources provided by the Stanford Genetics Bioinformatics Service Center. PMVECs from PAH and control patients were provided by the Pulmonary Hypertension Breakthrough Initiative, funded by the NIH and managed at University of Pittsburg. The tissues were procured at the Transplant Procurement Centers at Stanford University, managed by Marlene Rabinovitch and Roham T. Zamanian. The authors thank all patients and their proxies who participated in this study. The authors are also grateful to Patricia Angeles del Rosario (Stanford University Medical Center) for helping with the de-identified patient database.

## Source of funding

This work was supported by the National Institutes of Health (1R01HL17244901A1, 5R01HL13966407) to VA. de Jesus Perez, American Heart Association Postdoctoral fellowship (909301) and Career Development Award grant (24CDA1268257), Translational Research and Applied Medicine grant to S. Agarwal.

## Disclosures

V.A. de Jesus Perez reports support for the present manuscript from the National Institutes of Health National Heart, Lung, and Blood Institute. All other authors have nothing to disclose.

## SUPPLEMENTAL MATERIAL

Detailed methods

Supplementary Figures 1-5

Supplementary Figure legends 1-5

Supplementary tables 1-3

## NON-STANDARD ABBREVIATIONS AND ACRONYMS

PMVECs: pulmonary microvascular endothelial cells
BMPR2: bone morphogenetic protein receptor 2
CES1: carboxylesterase 1
pCES1: plasmid for CES1 overexpression
ROS: reactive oxygen species
FFA: free fatty acids
MDA: malondialdehyde
TG: triacylglycerol/triglyceride
scRNA: single cell RNA sequencing
RV: right ventricle
LV: left ventricle
cKO: conditional knockout
ECKO: endothelial specific CES1 cKO
HET-KO: heterozygous knockout
FAO: fatty acid oxidation
ChIP: chromatin immunoprecipitation
DE: differential expression
BP: biological processes
CC: cellular component
GO: gene ontology
GSEA: gene set enrichment analysis
NES: normalized enrichment score

## Notes

### Competing Interest Statement

The authors have declared no competing interest.

